# *Aimea* gen. nov. defines a novel plant-associated yeast genus in *Microbotryomycetes* with three novel species

**DOI:** 10.64898/2026.04.08.717246

**Authors:** Julian A. Liber, Marco A. Coelho, Sheng-Yang He

## Abstract

Plant tissues and surfaces are among the largest microbial habitats on Earth, and commensal yeasts are common members of these communities, where they can contribute to plant-microbe interactions including the biological control of plant diseases. Here, we describe a novel genus, *Aimea*, of unpigmented, plant-associated basidiomycete yeasts, in the class Microbotryomycetes, and name three new species (*A. erigeronia*, *A. cardamina*, and *A. sorghi*) represented by four isolates from leaves and roots of multiple hosts. We characterize these taxa through analyses of metabolic requirements, tolerance to differences in osmolarity, pH, and temperature, and enzymatic activities. In parallel, we generate near-chromosome-scale hybrid genomes annotated with transcriptome data. We employ whole-genome and multilocus phylogenetic approaches to infer the placement of these species within a monophyletic clade. We use comparative genomics to examine how the gene content of these yeasts differs from that of other members of the Microbotryomycetes, including an apparent proliferation of retrotransposons. We further demonstrate the genetic transformability of these taxa using *Agrobacterium tumefaciens*-mediated transformation. The description of these new species, together with high-quality genome resources and a genetic transformation protocol, establishes a foundation for experimental studies of these novel plant-associated yeasts and their interactions with hosts and other microbes.

## Introduction

Plant leaves and roots harbor diverse yeast communities that are increasingly recognized as important components of the plant microbiome (Glushakova and Chernov 2010; Boekhout et al. 2022; Sepúlveda et al. 2023). Among these are basidiomycetous yeasts within the class Microbotryomycetes (Pucciniomycotina) including *Microbotryum, Rhodotorula, Sporobolomyces, Rhodosporidiobolus, Leucosporidium, Mastigobasidium, Meredithblackwellia*, *Oberwinklerozyma*, *Microbotryozyma*, *Chrysozyma*, *Colacogloea*, *Heitmania, Slooffia*, *Pseudohyphozyma*, *Fellozyma*, and *Curvibasidium* (Li et al. 2020). Despite growing interest in the application of these yeasts for agricultural (Li et al. 2016) and industrial (Coradetti et al. 2018) uses, continued isolation of previously undescribed taxa indicates that their diversity and biology remain far from fully resolved (Jiang et al. 2024).

Microbotryomycete yeasts are also of ecological and applied interest. This group includes both plant pathogens, such as *Microbotryum* spp. (Hood et al. 2010), and species used for biocontrol of plant pathogens, such as *Rhodotorula* spp. (Castoria et al. 2005). Some members have been investigated for biotechnological applications, including lipid production (Coradetti et al. 2018), and genetic tools have been developed for several taxa (Takahashi et al. 2014; Pi et al. 2018; Jiao et al. 2019). They can be abundant in the phylloplane of some crops; for example, 27% of yeast colonies recovered from the romaine lettuce phylloplane were identified as a single *Sporobolomyces* species (*Sporobolomyces lactucae*) (Haelewaters et al. 2021; Fatemi et al. 2022).

While the Sporidiobolales (*Rhodotorula*, *Rhodosporidiobolus*, and *Sporobolomyces*) are distinctively pigmented, most other taxa within the class are not readily distinguished by pigmentation or morphology alone. Earlier classifications relied in part on traits such as cell wall composition and coenzyme Q (CoQ) systems (C. Kurtzman et al. 2011). Molecular phylogenetic approaches, often integrated with physiological and metabolic data (Li et al. 2020; Jiang et al. 2024), are now central to species and genus delimitation in this group. At the same time, expanding genomic resources from initiatives such as the Joint Genome Institute’s Fungal Genomics Program (Grigoriev et al. 2011) have improved phylogenetic resolution (Li et al. 2021) and enabled comparisons of gene content and function relevant to understanding ecological interactions (Martino et al. 2018).

Genetic transformation in basidiomycete yeasts has been pursued with four main approaches: *Agrobacterium tumefaciens*-mediated transformation (ATMT) (Lin et al. 2014), electroporation (Takahashi et al. 2014; Pi et al. 2018), protoplasting (Tully and Gilbert 1985), and gene gun/biolistic (Abbott et al. 2013) methods. Among these, ATMT in fungi, first described by Bundock et al. (1995), has been successfully applied across a broad range of fungi, including ascomycetes (Michielse et al. 2005; Petrucco et al. 2024), mucoromycetes (Ando et al. 2009), chytrids (Medina et al. 2020), and basidiomycetes. Unlike homology-based methods, the same construct can often be used across multiple strains and/or species, facilitating rapid protocol development. Furthermore, cells do not need to be made competent in the same manner as for electroporation (Pi et al. 2018), which further simplifies the procedure. T-DNA insertion is random (Tzfira et al. 2004) and construct expression may thus vary substantially depending on genomic context.

Here, we propose a novel yeast genus to accommodate three novel species isolated from leaves and roots, supported by molecular phylogenetic and metabolic analyses that define a previously undescribed monophyletic clade. We also generate near-chromosome-scale genome assemblies, annotated with transcriptome data, and demonstrate successful genetic transformation and heterologous gene expression via ATMT in two of the novel species.

## Methods

### Isolation and culturing

Yeast strains were isolated by ballistospore capture (Nakase & Takashima, 1993) or by planting ground leaf tissue. For ballistospore capture, a leaf was taped adaxial side-up onto the lid of a 100 mm 1X potato dextrose agar plate (PDA: 39 g/L potato dextrose agar, Difco, supplemented with 34 μg/mL chloramphenicol and 100 μg/mL ampicillin), and placing the agar-containing base over the leaf. The plates were kept in this orientation, sealed with Parafilm, and incubated at room temperature, before subculturing on PDA with antibiotics. Yeasts were additionally isolated by harvesting whole leaves, and grinding 1 leaf in 1 mL of 10 mM MgCl_2_ for 3 minutes at 30 Hz in a TissueLyser II (Qiagen) using three ZiO 3 mm beads, then plating 100 μL of undiluted leaf suspension, 10-fold, and 100-fold diluted on PDA1/2 +YE (19.5 g/L Difco potato dextrose agar, 7.5 g/L Bacto agar, and 1 g/L yeast extract, with 34 mg/L chloramphenicol and 100 mg/L ampicillin). Plates were incubated at room temperature and colonies were picked once they appeared. *A. sorghi* NB124-2 was obtained from the surface-sterilized roots of hybrid *Sorghum*. Freezer stocks in 50% glycerol were prepared for yeast strains upon identification by ITS sequencing, and maintained at -80°C for long-term storage. Subsequent culturing of strains was performed on YPD (10 g/L yeast extract, 20 g/L peptone, 20 g/L D-glucose, with 15 g/L Bacto agar added for solid media), YM (3 g/L yeast extract, 3 g/L malt extract, 5 g/L peptone, 10 g/L D-glucose, 15-20 g/L Bacto agar), or MEA+YE (10 g/L malt extract, 1 g/L yeast extract, 15 g/L Bacto agar).

### PCR and ITS-LSU sequencing

Polymerase Chain Reaction of the internal transcribed spacer (ITS) region of the rRNA operon, including ITS1-5.8S-ITS2 and partial large subunit (LSU), was performed with Green GoTaq Master Mix (Promega cat. no. M712), using 333 nM of the primers ITS1f (5’-CTTGGTCATTTAGAGGAAGTAA-3’;Gardes & Bruns, 1993) and LR3 (5’-CCGTGTTTCAAGACGGG-3’;Vilgalys & Hester, 1990), with yeast cells added directly to 12 μL reactions with a pipette tip. Cycling parameters were: 95°C for 10 min, 35 cycles of 95°C/30 s, 57°C/30 s, 72°C/1 min, followed by a 5 min hold at 72°C. Amplification of additional LSU sequence was performed with LR0R (5’-ACCCGCTGAACTTAAGC-3’) and LR7 (5’-TACTACCACCAAGATCT-3’), using 54.3°C as the annealing temperature. PCR products were visualized via gel electrophoresis on a 0.9% agarose gel, run for 12 minutes at 150 V and stained with SYBR Safe. PCR products were cleaned of primers and dNTPs with 0.24 U thermolabile exonuclease I (NEB cat. no. M0568) and 0.024 U QuickCIP (NEB cat. no. M0525) in 0.042x rCutSmart buffer and 0.043x NEBuffer r3.1 buffer, with 1.5 μL of PCR product in a 2.7 μL reaction. This mix was heated to 37°C for 15 minutes, then to 80°C for 1 minute. Cleaned PCR products were sequenced by Eurofins Genomics using the ITS1f primer for the ITS region and both LR0R and LR7 for the LSU region. ITS sequences (spanning ITS1f-LR3, including partial SSU, ITS1-5.8S-ITS2, and partial LSU) were searched against GenBank nucleotide and GlobalFungi (Větrovský et al. 2020) databases to attempt identification and determine the frequency of detection or isolation of related strains.

### Genome and transcriptome sequencing

Genomic DNA for whole-genome sequencing was extracted using the Wizard Genomic Purification Kit (Promega cat. no. A1120), modified by omitting enzymatic cell wall degradation and instead lysing cells at 65°C for 1 h in Nuclei Lysis Solution. For isolates *A. erigeronia* JL201 and *A. cardamina* JL221, 650 Mbp of short-read sequencing was generated by SeqCenter (Pittsburgh, PA, USA) on the Illumina NextSeq 2000 platform using paired-end 2 × 151 bp chemistry. Additional long-read sequencing was performed for all three species using Oxford Nanopore Technologies (ONT) v14 library preparation and R10.4.1 flow cells with an expected output of 360 Mb of sequence generated for each by Plasmidsaurus (Eugene, OR, USA). *A. sorghi* NB124-2 was further sequenced by Plasmidsaurus on the Illumina NextSeq 2000 platform with 2 × 151 bp chemistry, producing 113.6 Mbp of output. A transformed line of *A. erigeronia* JL201 containing a targeted genomic insertion was also sequenced using ONT; reads aligning to the inserted sequence with Minimap2 were removed, then pooled with reads from the wild-type strain prior to assembly.

RNA-Seq was performed on *A. erigeronia* JL201 to examine transcriptional responses to a variety of conditions, including salt stress (5% w/v NaCl), osmotic stress (20% w/v sucrose), pH stress (pH 3.5 or 7), redox stress (1.5 mM H_2_O_2_), nitrogen deprivation (0% ammonium sulfate), and carbon source shift (10 mM citrate and 0 % D-glucose), each compared to a standard synthetic defined medium (SD) containing 1x yeast nitrogen base without amino acids or ammonium sulfate (Millipore-Sigma cat. no. Y1251, 1.7 g/L), 0.5% ammonium sulfate, and 2% D-glucose, pH ∼4.8.

Briefly, a starter culture of 10 mL YPD in a 50 mL conical tube was inoculated from a single colony of *A. erigeronia* JL201 and grown for 2 days at 28°C and shaking at 225 rpm. The culture was diluted to 1050 mL of SD medium and aliquoted in 10 mL volumes into 15 mL conical tubes. These tubes were incubated as before, tilted slightly. After 48 h, the tubes were centrifuged at 3214 × g for 5 min at room temperature, then the T=0 h samples were snap frozen in liquid nitrogen. For the 1 h and 24 h time points, cells were transferred to 2 mL microcentrifuge tubes, then 1.5 mL treatment or control medium was added. After 1 h and 24 h of incubation in the same conditions, the cultures were centrifuged at 15000 × g for 1 min, the supernatant removed, and 450 μL of Q iagen RLT buffer with 1% ꞵ-mercaptoethanol was added before snap freezing. RN A was extracted using a hot acid phenol protocol (Collart and Oliviero 1993), treated with TURBO DNA-free DNase, and sequenced on an Illumina NovaSeq 6000 S4 lane with 2 × 150 bp format following polyA enrichment. The resulting sequencing reads allowed for annotation of gene positions.

### Genome assembly, annotation, and comparative genomic analyses

Genome and transcriptome data are available under BioProject PRJNA892096. Illumina sequencing reads were quality filtered with fastp (Chen et al. 2018). Long reads were assembled with Flye v2.9.2 (Kolmogorov et al. 2019), and the resulting assemblies were then polished with Polypolish v0.6.0 (Bouras et al. 2024) using the Illumina reads aligned with BWA-MEM (Li 2013). Gene models for JL201 were predicted using BRAKER3 v3.0.6 (Gabriel et al. 2024) implementing AUGUSTUS (Stanke et al. 2006) and GeneMark-ETP (Brůna et al. 2024) on the EU Galaxy platform (Jalili et al. 2020) integrating subsampled mRNA reads following alignment to the genome with HISAT2 v2.1.0 (Kim et al. 2019). The protein set predicted for JL201 was then used as evidence for gene prediction in the remaining genomes. Predicted proteins were functionally annotated with InterProScan (v5.59-91.0), implementing TIGRFAM, FunFam, SFLD, SUPERFAMILY, PANTHER, HAMAP, PROSITE, SMART, PRINTS, PIPSR, PROSITE, AntiFam, MobiDBLite, PIRSF, and UniProtKB.

Comparative genomic analyses based on InterProScan annotations generated for the three *Aimea* species and for proteomes of 12 other Microbotryomycetes taxa (Table S1). The relative abundance of annotation terms in *Aimea* species compared the other Microbotryomycetes, as well as among *Aimea* species, was evaluated using row-wise Fisher’s tests, implemented in rstatix v0.7.2 (Kassambara 2023), with *P* values adjusted with the Holm-Bonferroni method (Holm 1979).

Linear synteny plots of full contigs and mating-type (*MAT*) loci regions were generated with EasyFig v2.2.2 (Sullivan et al. 2011) using BLASTn for alignment. *MAT* loci were first identified by BLAST searches using previously characterized Microbotryomycetes *MAT* genes and their flanking genes as queries (Coelho et al. 2010; Maia et al. 2015). The *HD* loci were defined as the regions spanning the *HD1* and *HD2* genes. The *P/R* loci were preliminary delineated based on structural comparisons between opposite mating types from different species, rather than between opposite mating-type strains of the same species, because additional strains were not available. Accordingly, the inferred boundaries and gene content of the P/R loci should be regarded as provisional.

### Phylogenomic and multilocus phylogenetic analyses

Phylogenetic relationships within the Microbotryomycetes were inferred using a two-step approach in which a genome-scale ASTRAL species tree, derived from whole-genome data, was used to constrain a seven-locus maximum-likelihood phylogeny inferred with IQ-Tree.

Whole-genome shotgun assemblies were downloaded from GenBank and JGI Mycocosm, then BUSCO v6.0.0 (Tegenfeldt et al. 2025) was run using the Microbotryomycetes_odb12 profiles on these and the generated genome assemblies to extract nucleotide sequences. MAFFT v7.543 was then used to align nucleotide sequences, followed by gene-tree inference with IQ-Tree v2.2.2.7 (Minh et al. 2020). The species tree was then inferred using weighted ASTRAL v1.23.3.7 (Zhang et al. 2025).

Because most prior studies of the taxonomy of this group (Wang et al. 2015; Li et al. 2020; Jiang et al. 2024) used multi-locus data from rDNA (SSU, LSU, 5.8S) and four protein-coding loci (CYTB, TEF1, RPB1, and RPB2), these loci from type or reference genome strains in the Microbotryomycetes, Agaricostilbomycetes, Spiculogloeomycetes, Mixiomycetes, and Atractiellomycetes were incorporated into the species tree. Sequences from genomes were found using HMMs with HMMER v3.3.2 (Eddy 2011) or ITSx v1.1.3 (Bengtsson-Palme et al. 2013) trained on known sequences. Accessions and genomic coordinates are in Table S2. Each rRNA region and protein-coding locus was aligned using MAFFT, corrected and trimmed manually, and assigned to partitions. Each codon position for each protein-coding gene was assigned to a partition. ITS1 and ITS2 loci as well as intronic sequence from protein-coding loci were excluded from phylogenetic analysis due to poor alignment quality.

ModelFinder (Kalyaanamoorthy et al. 2017) implemented in IQ-Tree v2.2.2.7 (Minh et al. 2020) was used to select the best substitution model for each partition, then tree-inference was performed using the IQ-Tree algorithm followed by 1000 Ultra-Fast bootstraps (Minh et al. 2013). The Agaricostilbomycetes were designated as the outgroup. Trees were manipulated and displayed in FigTree v1.4 (Rambaut 2012) as with ggtree 3.8.2 (Yu et al. 2017). Gene trees for each locus were inferred using the same methodology.

### Yeast Phenotypic Characterization

Colony morphology was examined on YMA (5 g/L peptone, 3 g/L yeast extract, 3 g/L malt extract, 10 g/L D-glucose, 20 g/L agar) after 4 days of incubation at 25°C (Suh et al. 2008 Nov). Images were color corrected using a Calibrite color checker card in Adobe Photoshop. Cell morphology was examined by incubating cells in YMB (YMA without agar) for 5 days at 25°C.

Tests for carbon source assimilation and carbon source oxidation were performed with Biolog YT Microplates (Biolog cat. no. 1005), using cells grown on BUY agar plates (Biolog cat. no. 71005) for 2 days at 28°C, and resuspended in water to an optical density matching that of the YT standard (Biolog cat. no. 3415). Carbon assimilation and oxidation were assessed by absorbance at 590 nm (for assimilation) and 500 nm (for oxidation). The absorbance for each well was compared to the maximum absorbance for the assimilation or oxidation tests, with the negative control well set to zero. A substrate scored as positive for assimilation or oxidation for absorbances ≥50% of the maximum, and was counted as variable for absorbances ≥20% and <50% of the maximum. Substrates were counted as negative if the absorbance was <20% of the maximum (Haelewaters et al. 2020).

All other metabolic and physiological tests, including nitrogen assimilation, vitamin requirement, osmotic and chemical stress tolerance, and enzyme assays were performed according to Suh et al. (2008 Nov). Carbon sources were prepared at concentrations of 5 g/L in 6.7 g/L yeast nitrogen base without amino acids (BD cat. no. 291940), 45 mg/L yeast extract, and 45 mg/L casamino acids, and were sterilized using 0.22 μm filters. Soluble starch was autoclaved instead of filtered. ∼100 μL of methanol and 135 μL of absolute ethanol were added to 4.5 mL of pre-filtered base medium for their respective assimilation tests. DL-lactate and citrate were added as 5g/L of their conjugate acids, and their pH was adjusted to 5.0 with NaOH. D-lactose was additionally tested for *A. erigeronia* and *A. cardamina*. A positive control of D-glucose and a negative control without an added carbon source were included. Fermentation substrates were tested at 20 g/L in 10 g/L yeast extract. ∼18 mL of a given medium was added to a 20×150 mm glass test tube, containing an inverted 12×75mm test tube. The inner test tube was manipulated to remove any bubbles, then the tubes were capped and autoclaved for 15 minutes. Tested substrates included D-glucose, D-xylose, D-galactose, D-lactose, and D-sucrose with a negative control of no carbon source. *Saccharomyces cerevisiae* BY4741 was added to a D-glucose test tube to serve as a positive control. Both carbon assimilation and fermentation tests were inoculated by collecting cells of a 2 day-old YMA plate culture, grown at 25°C, suspending cells in water, and adding 100 μL of cell suspension per tube. Tubes were incubated at 25°C without shaking. Turbidity was observed for assimilation tests at 7 and 21 days. Fermentation tests were checked at 7, 14, and 21 to observe bubble formation.

Nitrogen assimilation assays were performed with the following base medium. 2x yeast carbon base was prepared with 20 g/L glucose, 40 μg/L biotin, 4 mg/L calcium pantothenate, 4 μg/L folic acid, 20 mg/L *myo*-inositol, 0.8 mg/L niacin, 0.4 mg/L *para*-aminobenzoic acid (PABA), 0.8 mg/L pyridoxine HCl, 0.4 mg/L riboflavin, 0.8 mg/L thiamine HCl, 2 mg/L L-histidine, 4 mg/L L-methionine, 4 mg/L L-tryptophan, and the following salts, per anhydrous amount: 1 mg/L H_3_BO_3_, 80 μg/L CuSO_4_, 0.2 mg/L KI, 0.4 mg/L FeCl_3_, 0.8 mg/L MnSO_4_, 0.4 mg/L Na_2_MoO_4_, 0.8 mg/L ZnSO_4_, 1.7 g/L KH_2_PO_4,_, 0.3 g/L K_2_HPO_4_, 1 g/L MgSO_4_, 0.2 g/L NaCl, 0.2 g/L CaCl_2_. Nitrogen sources were prepared separately at the following concentrations in 2x yeast carbon base: 0.6 g/L KNO_3_, 0.84 g/L NaNO_2_, 1.33 g/L L-lysine HCl, 1.08 g/L cadaverine HCl, 0.8 g/L creatine monohydrate, 0.68 g/L creatinine, 1.33 g/L D-glucosamine HCl, 0.40 g/L imidazole, and 1.28 g/L D-tryptophan. pH was adjusted to 5.5-6.5 with 3.7% HCl if needed, then 20g/L (final concentration) Bacto agar was added and an equal amount of water to 2x yeast carbon base. The media were then autoclaved and poured into 100 x 15 mm plates.

Vitamin requirement tests were performed using a base medium of 10 g/L D-glucose, 5 g/L Difco vitamin-free casamino acids, 1 g/L KH_2_PO_4_, 0.5 g/L MgSO_4_-7H_2_O, 0.1 g/L CaCl_2_-2H_2_O, and 0.1 g/L NaCl. This base solution was autoclaved, then filter-sterilized vitamin solutions were added at 10x concentration to achieve the same final concentrations as in the 1x nitrogen assimilation medium. These vitamins were biotin, calcium pantothenate, folic acid, *myo*-inositol, niacin, PABA, pyridoxine HCl, riboflavin, and thiamine HCl. Dropout medium without *myo*-inositol, calcium pantothenate, biotin, niacin, pyridoxine HCl, thiamine HCl, or PABA, singularly, and without biotin and thiamine HCl or pyridoxine HCl and thiamine HCl, doubly, were prepared. Cells from a YMA plate grown at 25°C for 2 days were suspended in the base medium for 5 days at 25°C to starve them of vitamins, before adding 100 μL to each culture tube with 5 mL of base medium with all vitamins, dropout medium missing 1 or 2 vitamins, or a negative control without any vitamins. Turbidity was assessed at 3, 7, and 14 days post inoculation.

Tolerance tests were performed on solid medium: acetic acid tolerance (1% glacial acetic acid added to 100 g/L D-glucose, 10 g/L tryptone, 10 g/L yeast extract, 20 g/L Bacto agar after autoclaving and cooling to 60°C), and the following osmolytes in 1% yeast extract and 2% Bacto agar medium: 50% (w/v) D-glucose, 60% (w/v) D-glucose, 10% (w/v) NaCl, 16% (w/v) NaCl, and a control with no osmolyte. A 10-fold dilution series from 1 to 10^6^ dilution of yeast cells grown on YMA plates for 3 days at 25°C and suspended in sterile water was prepared, and 10 μL drops were added to each of the tolerance media. Tolerance tests of high temperatures (30°C and 35°C) and cycloheximide (0.1% and 0.01%, at 25°C) were inoculated using the same cell suspension, and each was done in a solution of 6.7 g/L yeast nitrogen base without amino acids (BD cat. no. 291940) and 5 g/L D-glucose. Growth for tolerance tests was observed at 8 days post inoculation. pH range tolerance was tested using 1.7 g/L yeast nitrogen base without amino acid and ammonium sulfate (Millipore-Sigma cat. no. Y1251) with 5 g/L ammonium sulfate and pH adjusted to 3, 4, 6, 8, or 9 with hydrochloric acid or sodium hydroxide solutions prior to filter sterilization. Growth was assessed in 200 μL volumes in a 96-well microplate, after 4 days of incubation at 28°C.

Urease activity was tested in Urea R Broth (Rustigian and Stuart 1941), composed of: 20 g/L urea, 0.1 g/L yeast extract, 0.01 g/L phenol red, 0.095 g/L Na_2_HPO_4_, 0.091 g/L KH_2_PO_4_, with pH adjusted to 6.9 and filter sterilized. A loop of cells from a YMA plate grown at 25°C for 2 days was added to 0.5 mL of broth, which was then incubated at 37°C for up to 4 h or until color change compared to uninoculated broth was observed. A bright pink or red color indicated urease activity.

Gelatin liquefaction was assessed in a gelatin medium composed of 20% gelatin, 5 g/L D-glucose, and 6.7 g/L yeast nitrogen based. The medium was autoclaved and 2.5 mL was dispensed into culture tubes. Cells from a YMA plate grown at 25°C for 2 days were picked up with a long wood streaker and stabbed into the solidified medium. The culture tubes were then incubated at 25°C for 3 weeks. An uninoculated tube was simultaneously incubated for comparison.

Ballistospore formation was assessed by streaking each culture on a 100 x 15 mm corn meal agar (CMA; Criterion Dehydrated Culture Media, Hardy Diagnostics, Santa Maria, CA, USA) plate, and placing another CMA plate with a sterilized microscope slide below the streaked plate, then sealing the two plates with surgical tape (C.P. Kurtzman et al. 2011). These plates were incubated at room temperature, approximately 22°C, for 3 weeks, and examined for colony formation on the unstreaked plate. Crosses between JL221 and JL257 were attempted on V8, PDA, and YMA media by co-inoculating a light lawn of cells and incubating at room temperature in the dark up to 6 months.

### Genetic Transformation via *Agrobacterium tumefaciens*

Genetic transformation of yeast isolates was performed using an ATMT approach modified from Bundock et al. (1995). *Agrobacterium tumefaciens* AGL-1 was transformed via electroporation (Wolfram Jwd Debler 2017; Wolfram Jwd Debler 2019) with a binary vector plasmid (pMBM51) carrying a aphA1 (NEO) cassette for G418 resistance and enhanced yellow fluorescent protein (eYFP), both driven by the bidirectional histone H3-H4 promoter from *A. cardamina* JL221. This plasmid was originally derived from NM9-SpCas9-NLS3 (Schultz et al. 2019). NM9-SpCas9-NLS3 was a gift from Huimin Zhao (Addgene plasmid # 128177).

Yeasts were grown for 2 days in YPD at 28°C with shaking at 230 rpm in 10 mL culture tubes, then washed with water, diluted to an OD600 = 0.6. Agrobacterium cells were grown for 2 days in LB supplemented with kanamycin (50 mg/L) and rifampicin (20 mg/L) under the same conditions, then washed and diluted to an OD600 = 2.0-6.0. For co-cultivation, 50 μL each of yeast and Agrobacterium cells were added to each well of a 12-well microtiter plate containing 2 mL of 2% agar medium formulated in Bundock et al. (1995) at pH 5.2, with 100 μM acetosyringone, and three 3 mm glass spreading beads. Plates were gently agitated to spread the cells, then allowed to dry. The plates were sealed and incubated at 25°C for 3 days. After incubation, cells were resuspended in 300 μL of sterile water and plated on YPD containing 100 mg/L G418, 50 mg/L tetracycline, and 34 mg/L chloramphenicol. Fluorescent colonies were picked after 3-5 days of incubation at 28°C. T-DNA insertion locations were mapped using the splinkerette PCR method (Potter and Luo 2010) using BstYI digestion and confirmed by PCR across the insertion region.

## Results

### Yeast Isolation and preliminary identification

Three isolates (JL201, JL221, and JL257) with similar colony morphology were independently isolated from the leaves of *Erigeron* sp. (JL201) and *Cardamine hirsuta* (JL221 and JL257) in Durham, NC, USA from August 2021 to January 2022. These isolates fell into two groups with identical ITS-LSU sequences: group 1 (GenBank accession OQ388269, here described as *A. erigeronia*) and group 2 (OQ388277 and OQ388309, *A. cardamina*). A fourth isolate was obtained from a separate survey conducted in Ross Township, Michigan, USA in July 2021, during which isolate NB124-2 was isolated as an endophyte of *Sorghum bicolor* roots (OQ434130, *A. sorghi*). Additional attempts to isolate related strains were not successful so far. BLASTn searches of the NCBI GenBank Nucleotide Collection using each ITS-LSU sequence as a query did not identify additional deposited sequences, cultured or uncultured, with >90% identity as of March 27, 2026. Subsequent whole-genome sequencing, phylogenetic analyses, and physiological tests were performed on the initial representative isolate of each ITS-LSU group.

### Whole-genome sequencing and phylogenetic analyses support the recognition of three novel species

Genome assemblies were generated for strains JL201, JL221, and NB124-2 by combining Oxford Nanopore and Illumina sequencing.(Table 1). The final assemblies ofJL201, JL221, and NB124-2 are 17.591 Mbp, 19.439 Mbp, and 18.061 Mbp in length with N50 values of approximately 1.1-1.3 Mbp. Many contigs contained telomeric repeats at one or both ends (Table S3, Figure S1). BUSCO assessment of genome completeness using the Microbotryomycetes odb12 dataset recovered 92.5, 91.9, and 92.7% complete BUSCO genes in JL201, JL221, and NB124-2 assemblies, respectively. Analysis of GC content revealed differences across genomes, with JL221 showing a lower GC content (at 49.56%) than JL201 and NB124-2 (54.56% and 55.70%, respectively). Pairwise average nucleotide identity between the genomes as determined by OrthoANI (Lee et al. 2016) ranged from 69.13% to 72.44% (Table S4) indicating substantial genomic divergence among the three taxa and supporting their separation as distinct species (Lachance et al. 2020).

**Table 1.** Genome sequencing and assembly statistics.

| Statistic | *A. erigeronia* JL201 Hybrid | *A. cardamina* JL221 Hybrid | *A. sorghi* NB124-2 Hybrid |
| --- | --- | --- | --- |
| Main genome contig total | 32 | 21 | 25 |
| Main genome contig sequence total (MB) | 17.591 | 19.439 | 18.051 |
| Main genome gap percentage | 0.000% | 0.000% | 0.000% |
| Main genome contig N/L50 | 7/1.14 MB | 7/1.269 MB | 7/1.168 MB |
| Main genome contig N/L90 | 17/320.919 KB | 15/729.44 KB | 16/621.28 KB |
| Max contig length | 1.944 MB | 2.083 MB | 1.859 MB |
| Number of contigs > 50 KB | 24 | 18 | 21 |
| % main genome in contigs > 50 KB | 99.35% | 99.63% | 99.35% |
| % GC | 54.56 | 49.56 | 55.7 |
| Sequencing depth (bp) - Illumina | 1,481,457,092 | 696,692,023 | 113,613,450 |
| Coverage - Illumina | 84 | 36 | 6 |
| Sequencing depth (bp) - ONT | 442,168,009 | 1,971,258,087 | 1,204,826,042 |
| Coverage - ONT | 25.14 | 101.41 | 66.75 |
| Genes | 8459 | 7891 | 7830 |

### Phylogenetics

During the assembly of the phylogenetic dataset for the Microbotryomycetes, several publically available loci sequences were identified as likely contaminants because their best BLASTn matches corresponded to distantly related fungi, such as *Candida* species. These loci were excluded from the analysis (notably RPB1 from *Libkindia masarykiana* KT310 and *Yurkovia medeliana* KT152), but other loci from the same strains were retained if sequence similarity suggested that they were not contaminants. Additionally, one taxon, *Reniforma strues* (Pore and Sorenson 1990) was examined in greater detail because it showed both an unusually long branch in preliminary analyses and inconsistent placement of its sequenced loci when compared against type-strain sequences. Jiang et al. (2024) also observed long branch length and inconsistent placement of *R. strues*. To determine whether the publicly available sequences were all from the same organism and to verify its phylogenetic placement, we sequenced and assembled the genome of the type strain (NRRL Y-17275). The genome-derived sequences matched those previously deposited in GenBank, but their distance from described species suggested that *R. strues* is unlikely to belong within Microbotryomycetes. Because resolving the precise taxonomic placement of this taxon is outside the scope of the present study, it was excluded from downstream analyses.

The seven-locus genome-restrained phylogeny (Figure 1) of Microbotryomycetes places all three isolates in a fully supported monophyletic clade sister to the Microbotryales (including *Microbotryum, Microbotryozyma*, *Bauerozyma*, and *Ustilentyloma*) and *Yuzhangia*. The novel isolates and the Microbotryales have a high confidence placement in a clade with *Sampaiozyma*. Furthermore, these clades are inferred to be sister to *Leucosporidium*. A clade supported at 95% places the Sporiodiobolales (*Rhodotorula*, *Rhodosporidiobolus*, and *Sporobolomyces*) with *Curvibasidium*, *Pseudoleucosporidium*, *Nakaseozyma*, *Baiomyces*, and *Fengyania*. A fully-supported clade places the Sporiodiobolales and sister taxa with *Heitmania* and the clade containing *Leucosporidium*, *Sampaiozyma*, the Microbotryales, *Yuzhangia*, and the novel isolates. Because of the placement of all three isolates in a fully-supported monophyletic clade, with branch length similar to that of other genera in the order, we nominate these isolates as three novel species in the novel genus *Aimea*: *A. erigeronia* (JL201), *A. cardamina* (JL221 and JL257), and *A. sorghi* (NB124-2). As predicted by the average nucleotide diversity, *A. sorghi* and *A. erigeronia* with 72.4% ANI were inferred to form an exclusive clade, with *A. cardamina* (69.4 and 69.1% ANI to *A. sorghi* and *A. erigeronia*) as sister. The backbone of the Microbotryomycetes is composed of several short branches, many with low confidence, which preclude us from making durable family- and order-level taxonomic placements. However, the Microbotryomycetes are fully supported as a monophyletic clade.

**Figure 1.**
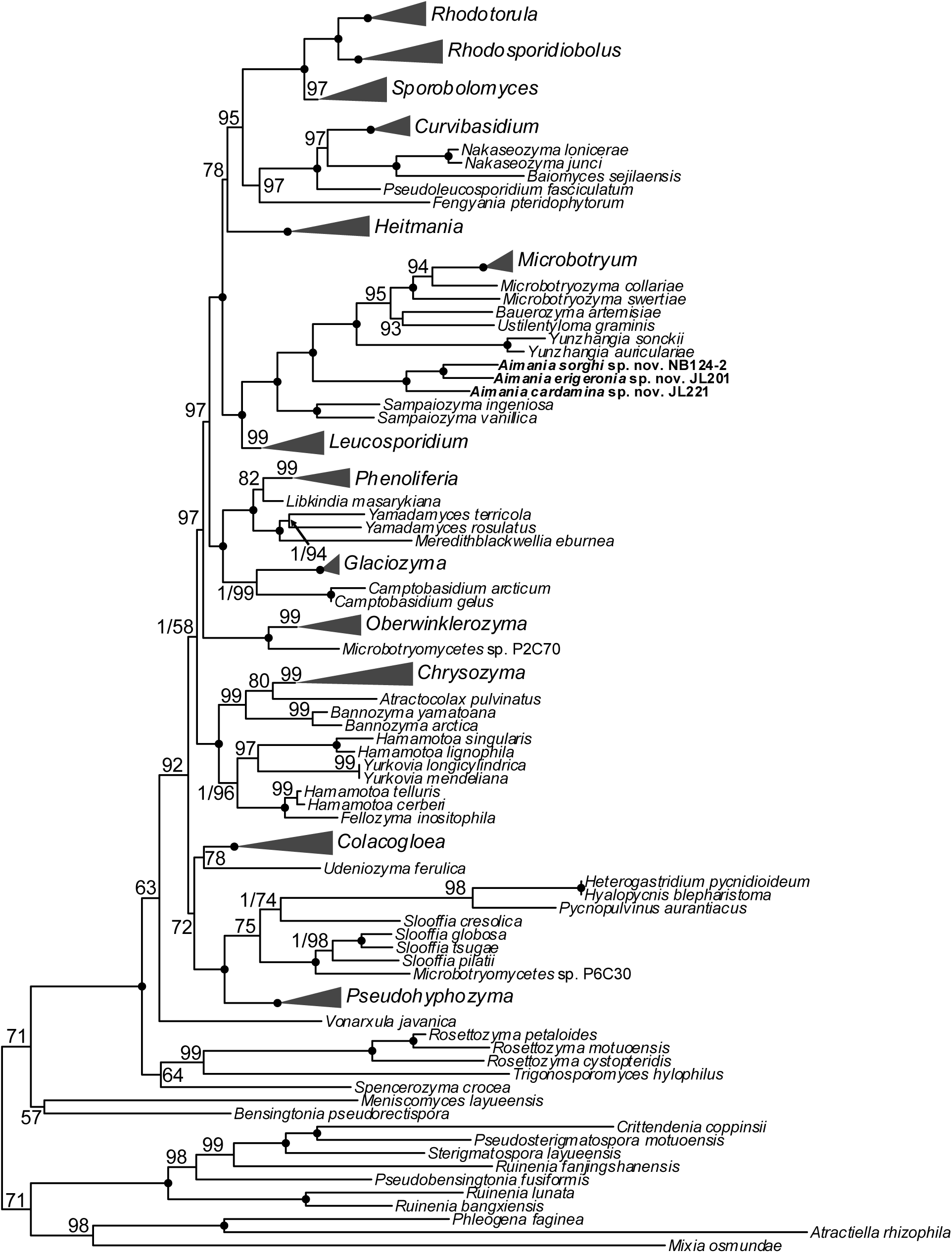
Multi-locus phylogeny of maximum likelihood phylogeny of Microbotryomycetes. Phylogeny of Microbotryomycetes inferred in two steps with an ASTRAL tree generated with whole genomes, then used to constrain a seven-locus (SSU, LSU, 5.8S, *CYTB*, *TEF1*, *RPB1*, and *RPB2*) maximum likelihood phylogeny generated with IQ-TREE. Node support is indicated by solid circles for 100% ultrafast bootstrapping (UFBoot2) support and 1.0 local posterior probability (LPP; if determined in ASTRAL). Otherwise, the UFBoot2 value and LPP are displayed (LPP/UFBoot2 or UFBoot2). Genera which were inferred as monophyletic clades are collapsed for display.

### Comparative Genomics

A comparative genomics approach was used to compare the predicted proteomes of *Aimea* spp. with predicted proteomes of other members of Microbotryomycetes (Figure 2), in the genera *Heterogastridium*, *Hyalopycnis*, *Leucosporidium*, *Meredithblackwellia*, *Microbotryum*, *Phenoliferia*, *Rhodotorula*, and *Sporidiobolus*, as well as Microbotryomycetes spp. P2C70 and P6C30 from JGI Mycocosm, which are likely *Oberwinklerozyma* sp. and *Slooffia* sp., respectively, given the inferred phylogeny. These proteomes were functionally annotated using the InterProScan pipeline for both *Aimea* and the publically-available datasets. However, genome assemblies and gene model annotations were performed independently and from variable types of sequencing data. In *Aimea* spp. the most strongly depleted annotation terms included ribonuclease inhibitors (and RNI-like proteins), proteins containing MYND-type zinc finger domains, furins, F-box-like proteins, and proteins with a proline rich extensin signature (Figure 2A). Conversely, proteins likely originating from transposons were the most enriched in *Aimea* spp., including integrases, reverse transcriptases, polyproteins, and DNA/RNA polymerases. These annotations were clustered in at most a single site per contig (Figure S1), suggesting that they may be localized to centromeres. When examining differences within the genus (Figure 2B), this enrichment of transposon components is most pronounced in *A. cardamina* JL221 (69 integrase and 108 reverse transcriptase Pfam domains), notably depleted in *A. sorghi* NB124-2 (8 and 55), and slightly depleted in *A. erigeronia* JL201 (27 and 51). The most notably metabolic annotations which differ are those for ATP citrate synthase/citrate lyase, which shows some enrichment in *A. cardamina* JL221.

**Figure 2.**
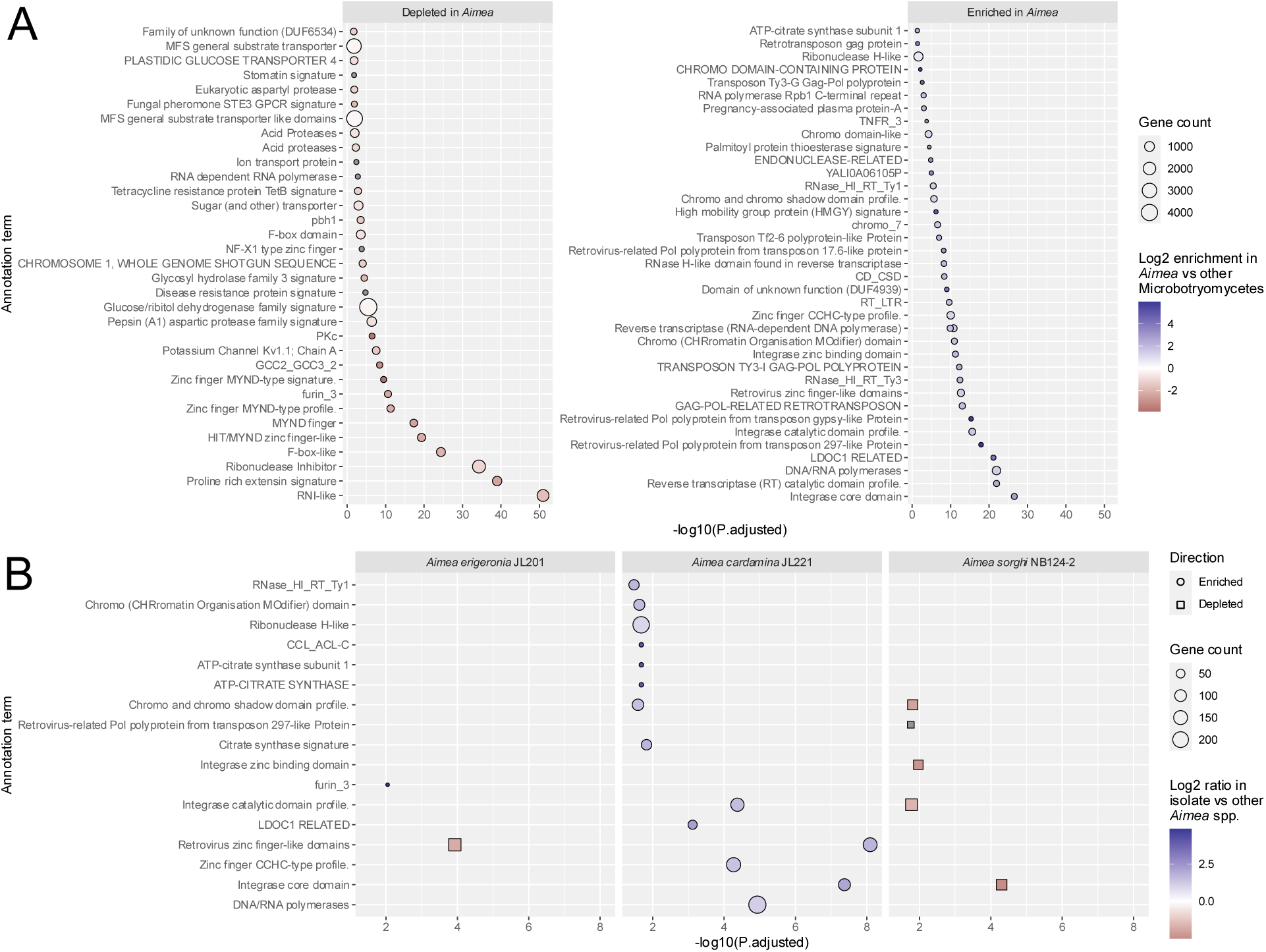
Enrichment of annotation terms in *Aimea* and its species. Annotation terms derived from InterProScan were compared in relative proportion between *Aimea* and twelve other members of the Microbotryomycetes (A) or between each species and the other two within *Aimea* (B). Significance was determined with Fisher’s exact test, and P values were adjusted within each analysis.

### Yeast Phenotypes

Yeast cultures streaked onto plates appeared generally similar, as cream colored, entire margin, and butyrous (Figure 3). Phenotypic characterization of the novel yeast species *A. erigeronia*, *A. cardamina*, and *A. sorghi* revealed that these yeasts share a very similar macro-and micromorphology, absence of hyphae or pseudohyphae, absence of fermentation of any tested carbon sources, generally similar carbon and nitrogen assimilation profiles, and similar tolerance to acetic acid, cycloheximide, and osmolytes, and share tolerated ranges of pH and temperature. However, notable differences in each species’ vitamin requirements and the ability to assimilate some specific carbon sources are observed (D-mannitol, L-malic acid, D-ribose, L-rhamnose, and Tween 80). The full set of analyzed traits is found in Table S5.

**Figure 3.**
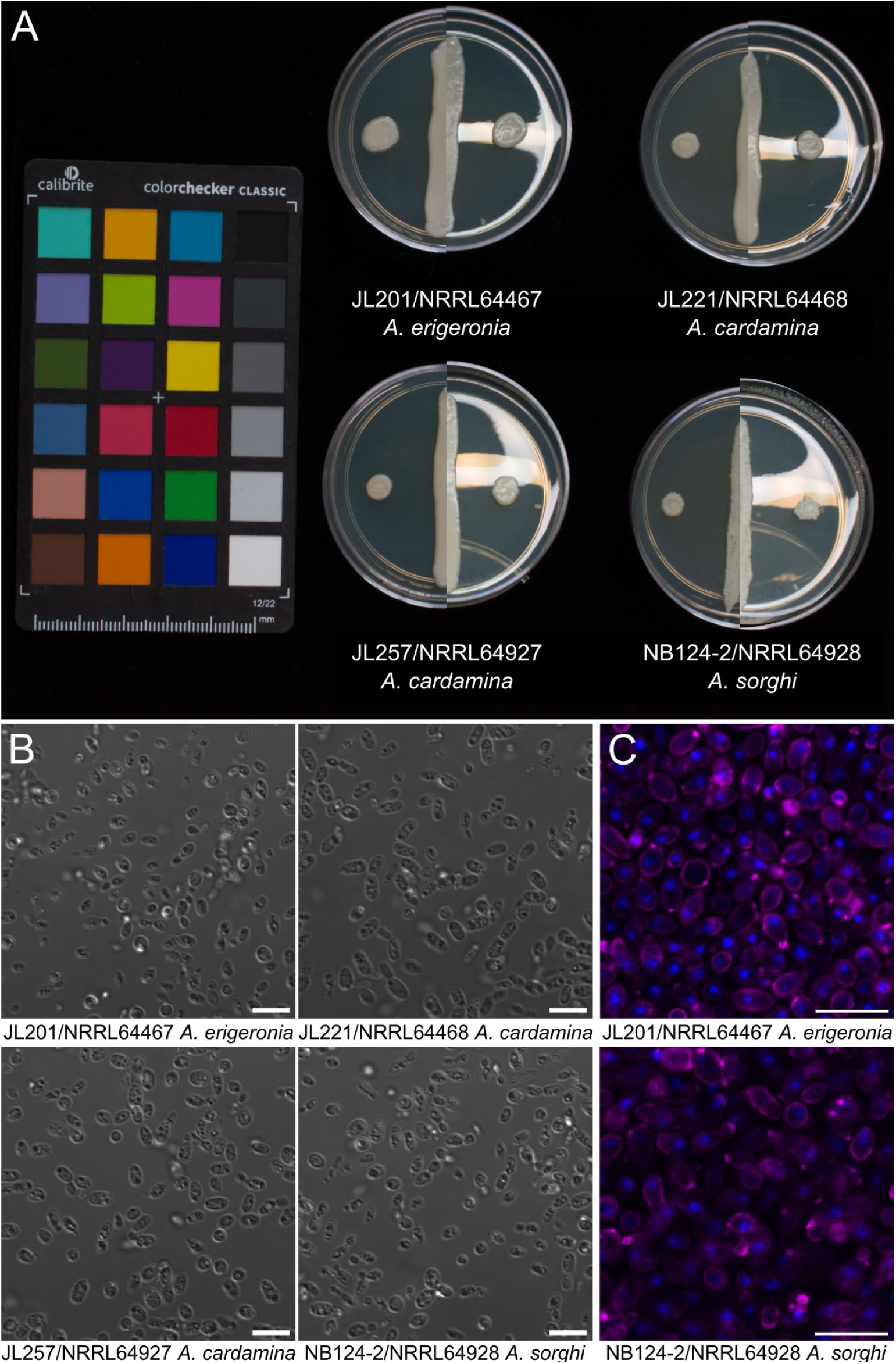
Macro- and micromorphology of *Aimea* strains. A) Composite of colony macromorphology after 4 days of incubation on YMA at 25°C, using both reflecting and non-reflecting lighting. B) DIC micrograph demonstrating cell micromorphology of planktonic cells grown in YMB for 5 days at 25°C. C) Cells stained with the DNA stain DAPI (blue) and the cell-wall binding stain wheat germ agglutinin (magenta; WGA-640R) observed with confocal laser scanning microscopy. B-C) Scale bars measure 10 μm.

### Genetic Transformation

Genetic transformation of two isolates of *Aimea* spp. is demonstrated by PCR and heterologous expression of eYFP (Figure 4). Each transformation produced 100s to 1000s of colony forming units, of which a minority showed strong expression of the fluorescent marker. Mapping of the insertion locations with splinkerette PCR was performed for first the right border of the T-DNA construct, then the sites were attempted to be confirmed by PCR amplification across the insertion locus. While this mapping was successful for *Aimea erigeronia* JL201, the absence of any band for *Aimea sorghi* NB124-2 led us to sequence from the left border using ONT linear amplicon sequencing. Of 2,931 reads, a single long read mapped outside the left border into contig 3 at position 527,250 (total contig length of 1,329,339 bp), a different large contig from the right border, which mapped to position 190,852 on contig 20 (total contig length of 329,703 bp). The mapping of reads to the middle of two large contigs suggest that a chromosome translocation occurred during the transformation, and the large number of reads extending into the plasmid sequence after the left border suggests that multiple copies of concatenated T-DNA were inserted into the genome.

**Figure 4.**
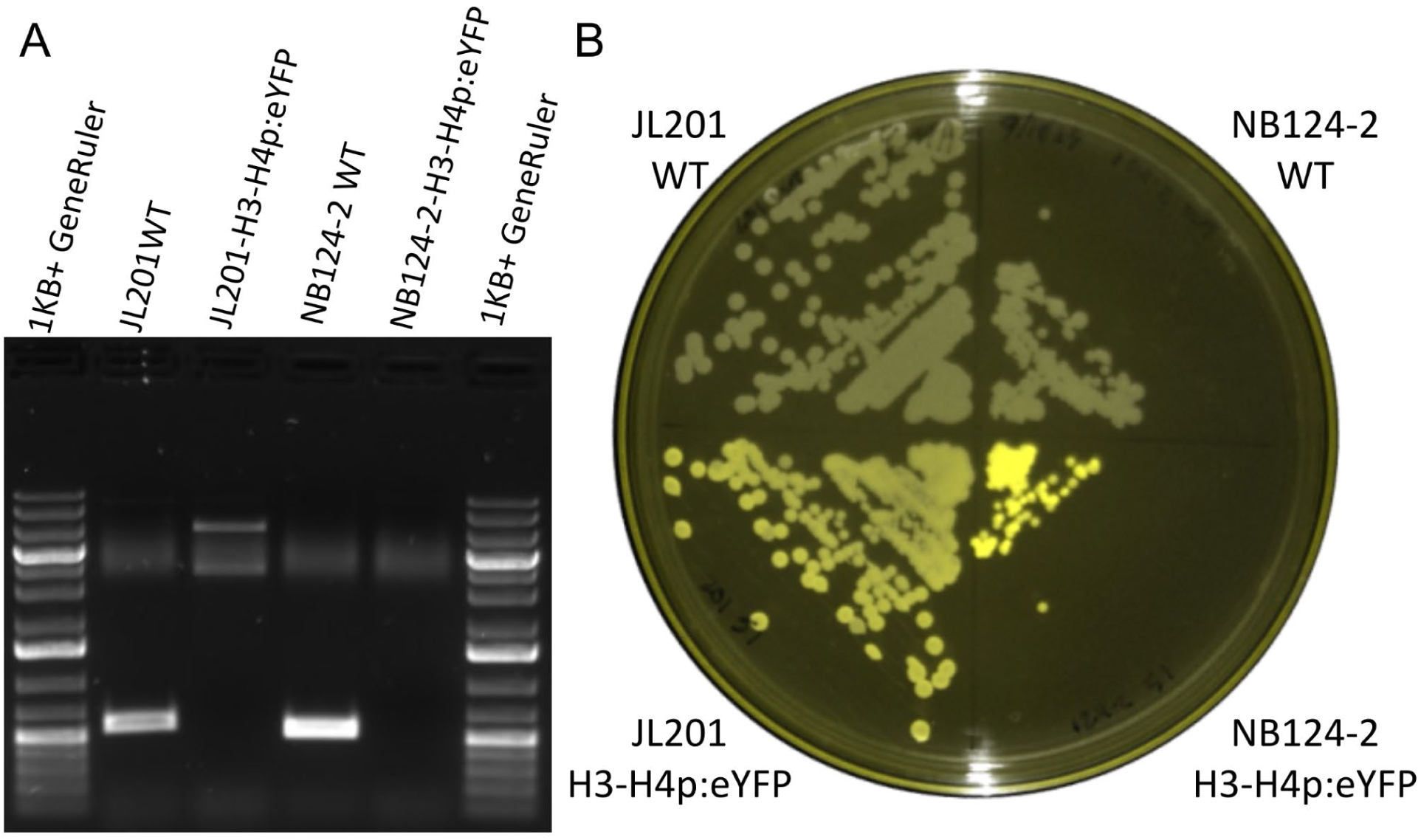
Transformation of strains JL201 and NB124-2. Isolates JL201 (*A. erigeronia*) and NB124-2 (*A. sorghi*) were genetically transformed to express eYFP via *Agrobacterium tumefaciens*. A) PCR using primers bridging mapped insertion sites for wild-type and transformed lines. The double bands in lane 3 in the JL201 transformant may be due to concatenated T-DNA insertions in this line, and possible induction of a local aneuploidy. The absence of the band in lane 5 in the NB124-2 transformant appears to be due to both concatenated T-DNA insertions at the locus and likely chromosomal translocation, evidenced by Sanger sequencing from the left and right borders. B) False-color fluorescence signal of transformed colonies, captured using an iBright™ FL1500 Imaging System with excitation filter range 455-485nm and emission range 508-537nm.

### Mating Locus

Mating compatibility in basidiomycetes is often governed by two *MAT* loci, the pheromone/receptor (*P/R*) locus and the homeodomain (*HD*) locus. When these loci are unlinked, the system is tetrapolar, whereas bipolar systems result when they become linked or when one locus no longer determines mating type (Fraser et al. 2007). For the three examined species, only *A. sorghi* has the *HD* and *P/R* loci on separate telomere-to-telomere contigs, supporting a tetrapolar mating system in which each locus segregates independently during meiosis (Figure 5). However, the near-chromosome-scale contigs of *A. cardamina* and *A. erigeronia*, together with their high degree of synteny with *A. sorghi*, also support the inference of a tetrapolar system in these two additional species.

**Figure 5.**
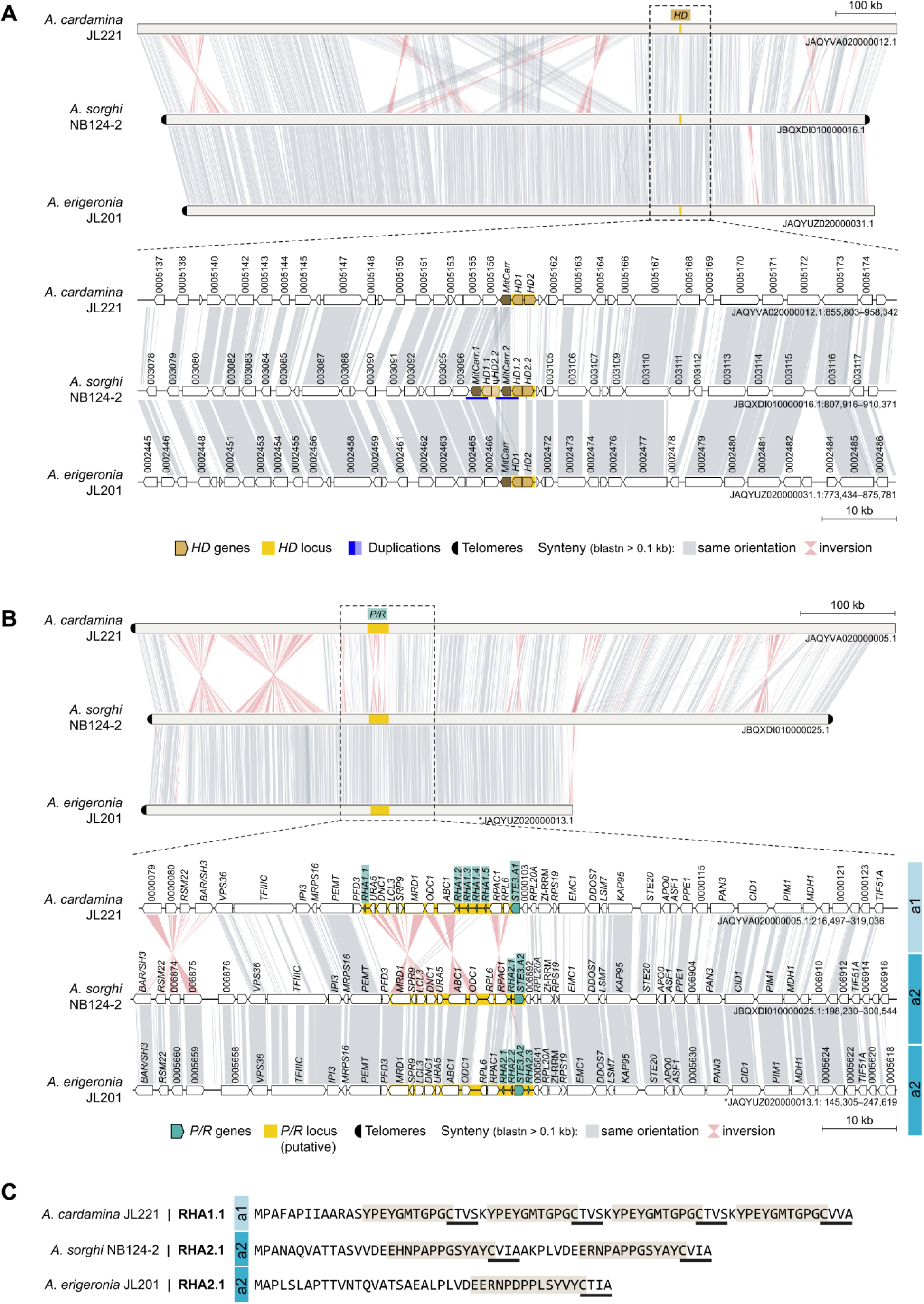
Organization and comparison of the *MAT* loci in three Aimea species. (A,B) Comparison of the *HD* locus (A) and the putative *P/R* locus (B) in three Aimea species. In each panel, the top subpanel shows a synteny view of the full contig, highlighting the chromosomal location of the focal locus, and the bottom subpanel provides a zoomed-in synteny view centered on that region. Conserved syntenic blocks are connected by shaded ribbons, with gray indicating the same orientation and pink indicating inversions. Locus-specific genes and the corresponding locus are highlighted as indicated in the key, together with other annotated features. In panel A, the *HD* region of *A. sorghi* contains two partially identical *HD1* copies that differ at their 5′ ends, consistent with a local duplication event (blue-highlighted region), and a partial *HD2* fragment fused to a neighboring flanking gene. Chromosomes inverted relative to their original assembly orientations are marked with asterisks. (C) Comparison of pheromone precursor proteins encoded in the *P/R* region. Repeated sequence segments proposed to correspond to the peptide moiety of the mature pheromone are shaded, and sequences resembling the C-terminal CAAX prenylation motif are underlined.

All three species encode an *HD* locus with broadly similar structure, composed of the typical divergently transcribed *HD1* and *HD2* homeodomain genes (Figure 5A). However, the *HD* region of *A. sorghi* includes two partially identical *HD1* copies that differ at their 5’ ends, consistent with a local duplication event, as well as a partial *HD2* fragment fused to a neighboring flanking gene. Aside from this feature, the *HD* locus sequences are more similar between *A. sorghi* and *A. erigeronia* than between either of those species and *A. cardamina*, consistent with their closer relatedness.

A similar pattern is also apparent at the *P/R* locus, where *A. sorghi* and *A. erigeronia* carry the *P/R a2* allele, whereas *A. cardamina* carries the opposite *P/R a1* allele, as inferred from sequence similarity of both the predicted receptor and mating pheromone (Coelho et al. 2008; Maia et al. 2015; Xu et al. 2016) (Figure 5B). The inferred *a1* and *a2 P/R* alleles also differ in gene order and orientation across the compared regions, with several inversions apparent in the synteny plots. Because these comparisons were performed among different species representing opposite inferred mating types, rather than opposite mating-type strains within each species, the precise boundaries and full gene content of the *P/R* locus in each species should be considered provisional. Despite this caveat, several genes within the predicted *P/R* locus regions of *Aimae* spp. were also identified in the *P/R* loci of both Sporidiobolales (Coelho et al. 2010; Liu et al. 2025) and Leucosporidiales (Maia et al. 2015), suggesting a shared evolutionary origin.

Variation within the *P/R* region was also evident among *A. sorghi* and *A. erigeronia*, which share the same inferred *a2* allele. *A. sorghi* contains a single pheromone gene (*RHA2*) that potentially encodes two peptide repeats, one differing from the other by a single amino acid, whereas *A. erigeronia* contains three identical pheromone genes, each encoding only a single copy of the predicted peptide moiety. By contrast, *A. cardamina* contains five identical pheromone genes (*RHA1*) at the *P/R a1* allele, each encoding four copies of the predicted peptide moiety. Despite evidence of a heterothallic mating system, the lack of additional known isolates of these species precludes experimental validation of the mating system.

#### Taxonomy

##### Aimea

Liber, Coelho & He, gen. nov.

MycoBank ###

Typification: *Aimea erigeronia* Liber, Coelho, & He.
Etymology: "Aimea", in honor of Dr. M. Cathie Aime, who is an accomplished mycologist, mentor, educator, and inspiration to a generation of mycologists.

This genus is proposed for the clade represented by JL201/NRRL 64467, which formed a distinct clade with JL221/NRRL 64468 and NB124-2/NRRL 64928 separate from other taxa in Microbotryomycetes (Figure 2). Sexual reproduction not observed. Colonies are cream-colored and butyrous. Cells produce polar buds from a narrow base, and are ovoid, ellipsoidal, or oblong. Hyphae or pseudohyphae not observed. Ballistoconidia are not observed in culture.

##### Aimea erigeronia

Liber, Coelho, & He, sp. nov.

MycoBank ###

Etymology: growing in/on *Erigeron* sp.; erigeronia = of Erigeron (fleabanes)
Typification: USA. NORTH CAROLINA: Durham County, Durham, Duke University West Campus, 36.00°N, 78.93°W, 100 m elevation, ballistospore isolate from *Erigeron* sp. leaf in vegetated strip along road, August 27, 2021. J.A. Liber (holotype NRRL 64467, preserved in a metabolically inactive state, ex-type CBS 18377).

Culture characteristics: Colonies from streak culture are cream colored, butyrous, smooth, and without pseudohyphae on YPD. Cells grown in YM broth for 5 days are ellipsoidal to ovoid, 5.34±0.61 μm long by 4.29±0.47 μm wide (mean ± standard deviation, n=51), with polar budding from a narrow base. No evidence of pseudohyphae in Dalmau culture. Absence of ballistospores after 3 weeks on CMA at 20-22°C.

Physiological and biochemical characteristics: Negative for fermentation of D-glucose, D-galactose, sucrose, lactose, raffinose, and D-xylose. Positive for assimilation of D-glucose, L-sorbose, D-glucosamine, sucrose, maltose, α,α-trehalose, methyl-α-D-glucoside, raffinose, melezitose, starch, glycerol, D-mannitol, 2-keto-D-gluconate, D-gluconate, succinate, citrate, palatinose, L-malate, and Tween 80 as sole carbon sources. Weak or delayed assimilation of D-galactose, D-xylose, D-arabinose, cellobiose, salicin, arbutin, inulin, xylitol, DL-lactate, and gentiobiose as sole carbon sources. D-ribose, L-arabinose, L-rhamnose, melibiose, lactose, erythritol, myo-inositol, methanol, and ethanol are not assimilated. Positive for assimilation of ammonium sulfate as a sole nitrogen source. Weak or delayed assimilation of L-lysine hydrochloride, cadaverine hydrochloride, glucosamine, and D-tryptophan as sole nitrogen sources. Potassium nitrate, sodium nitrite, creatine, creatinine, and imidazole are not assimilated. Growth in vitamin-free medium, but reduced growth in absence of thiamine hydrochloride. Positive for urease activity, negative for extracellular starch formation and production of acetic acid. Negative for growth with 1% acetic acid, 16% NaCl, or 60% (w/v) D-glucose (w/v), or 0.1% cycloheximide. Positive for growth in 10% NaCl, 50% (w/v) D-glucose, or 0.01% cycloheximide. Maximum growth temperature is 30°C. Reduced growth at pH=3. No growth at pH=9. Weak liquefaction of gelatin.

Physiologically, *A. erigeronia* differs from the closely related species *A. cardamina* and *A. sorghi* by its ability to assimilate D-mannitol, L-malic acid, and its ability to grow in vitamin-free medium. Morphologically, its cells are more rounded at a length:width ratio of 1.24.

##### Aimea cardamina

Liber, Coelho, & He, sp. nov.

MycoBank ###

Etymology: growing in/on *Cardamine hirsuta*; cardamina = of Cardamine (bittercresses)
Typification: USA. NORTH CAROLINA: Durham County, Durham, Duke University West Campus, 36.00°N, 78.94°W, 110 m elevation. Ground whole unsterilized leaf tissue of *Cardamine hirsuta* plated on PDA, from a grassy vegetated strip along a road. October 27, 2021. J.A. Liber (holotype NRRL 64468, preserved in a metabolically inactive state, ex-type CBS 18378). Other examined strains: JL257 (NRRL 64927, CBS 19403).

Culture characteristics: Colonies from streak culture are cream colored, butyrous, smooth, and without pseudohyphae on YPD. Cells grown in YM broth for 5 days are oblong to ovoid, 6.13±0.72μm long by 3.52±0.46 μm wide (mean ± standard deviation, n=51), with polar budding from a narrow base. No evidence of pseudohyphae in Dalmau culture. Absence of ballistospores after 3 weeks on CMA at 20-22°C.

Physiological and biochemical characteristics: Negative for fermentation of D-glucose, D-galactose, sucrose, lactose, raffinose, and D-xylose. Positive for assimilation of D-glucose, D-ribose, L-rhamnose, sucrose, α,α-trehalose, raffinose, starch, glycerol, 2-keto-D-gluconate, D-gluconate, succinate, citrate, palatinose as sole carbon sources. Weak or delayed assimilation of D-galactose, L-sorbose, D-glucosamine, D-xylose, L-arabinose, D-arabinose, maltose, methyl-α-D-glucoside, cellobiose, salicin, arbutin, melezitose, inulin, DL-lactate, and gentiobiose as sole carbon sources. Negative for assimilation of melibiose, lactose, erythritol, xylitol, D-mannitol, myo-inositol, methanol, ethanol, L-malate, and Tween 80. Positive for assimilation of ammonium sulfate and glucosamine as sole nitrogen sources. Weak or delayed assimilation of L-lysine hydrochloride, cadaverine hydrochloride, D-tryptophan. Potassium nitrate, sodium nitrite, creatine, creatinine, and imidazole are not assimilated. Growth in vitamin-free medium is negative, but is restored with addition of thiamine hydrochloride. Positive for urease activity, negative for extracellular starch formation and production of acetic acid. Negative for growth with 1% acetic acid, 16% (w/v) NaCl, or 60% (w/v) D-glucose (w/v), or 0.1% cycloheximide. Positive for growth in 10% (w/v) NaCl, 50% (w/v) D-glucose, or 0.01% cycloheximide (weak). Maximum growth temperature is 30°C. Reduced growth at pH=3. No growth at pH=9. Strong liquefaction of gelatin.

Physiologically, *A. cardamina* differs from the closely related species *A. erigeronia* and *A. sorghi* by its ability to assimilate L-rhamnose and its inability to grow in vitamin-free medium. Morphologically, its cells are more ellipsoidal than either species with a length:width ratio of 1.74.

##### Aimea sorghi

Liber, Coelho, & He, sp. nov.

MycoBank ###

Etymology: growing in/on *Sorghum bicolor*; sorghi = of Sorghum
Typification: USA. MICHIGAN: Kalamazoo County, Ross Township, Kellogg Biological Station, 42.39°N, 85.37°W, 285 m elevation. Ground surface-sterilized roots of hybrid *Sorghum bicolor* plated on PDA, from a grassy vegetated strip along a road. July 24, 2021. N. Beculheimer (holotype NRRL 64928, preserved in a metabolically inactive state, ex-type CBS 19404).

Culture characteristics: Colonies from streak culture are cream colored, butyrous, smooth, and without pseudohyphae on YPD. Cells grown in YM broth for 5 days are ellipsoidal to ovoid, 5.22±0.75 μm long by 3.60±0.55 μm wide (mean ± standard deviation, n=33), with polar budding from a narrow base. No evidence of pseudohyphae in Dalmau culture. Absence of ballistospores after 3 weeks on CMA at 20-22°C.

Physiological and biochemical characteristics: Negative for fermentation of D-glucose. Positive for assimilation of D-glucose, sucrose, maltose, α,α-trehalose, salicin, starch, citrate, and Tween 80 as sole carbon sources. Weak or delayed assimilation of D-galactose, L-sorbose, methyl-α-D-glucoside, raffinose, melezitose, glycerol, D-mannitol, 2-keto-D-gluconate, DL-lactate, palatinose, L-malate as sole carbon sources. Negative for assimilation of D-glucosamine, D-ribose, D-xylose, L-arabinose, D-arabinose, L-rhamnose, cellobiose, arbutin, melibiose, inulin, erythritol, xylitol, D-gluconate, methanol, ethanol, and gentiobiose as sole carbon sources. Positive for assimilation of ammonium sulfate, potassium nitrate, sodium nitrite, and cadaverine hydrochloride as sole nitrogen sources. Weak or delayed assimilation of L-lysine hydrochloride as a sole nitrogen source. Negative for assimilation of creatine, creatinine, glucosamine, imidazole, and D-tryptophan as sole nitrogen sources. Growth in vitamin-free medium is negative, but is restored with addition of thiamine hydrochloride. Positive for urease activity, negative for extracellular starch formation and production of acetic acid. Negative for growth with 1% acetic acid, 16% (w/v) NaCl, or 60% (w/v) D-glucose (w/v), or 0.1% cycloheximide. Positive for growth in 10% (w/v) NaCl, 50% (w/v) D-glucose (weak), or 0.01% cycloheximide. Maximum growth temperature is 30°C. Strong liquefaction of gelatin.

Physiologically, *A. sorghi* differs from the closely related species *A. erigeronia* and *A. cardamina* by its ability to assimilate nitrate, and nitrite, and its inability assimilate D-glucosamine, D-arabinose, or D-gluconate. Morphologically, its cells are intermediate of the other species with a length:width ratio of 1.43.

## Discussion

Here, we present the description of a novel basidiomycete yeast genus containing three species, all isolated from plant leaves and roots. These species are characterized for their morphology, metabolism, stress tolerance, genomic sequence and overlying gene models, and inferred phylogenetic relationships. The combination of these lines of evidence suggests that the four isolates presented represent three novel species, all within a novel monophyletic clade of similar divergence to other genus-level clades in the Microbotryomycetes.

The inferred phylogeny generally shows a similar structure to that inferred by Jiang et al. (2024), but some notable differences were observed. Several nodes that were poorly supported in previous studies (Wang et al. 2015; Li et al. 2020; Jiang et al. 2024) are more strongly resolved here. One of the clearest examples is the Microbotryales, which itself is well supported and also forms a well-supported clade with *Sampaiozyma* and the novel genus *Aimea*. Microbotryales was originally described to contain *Microbotryum* and *Ustilentyloma* (Bauer et al. 1997), which is now supported by both multi-locus and genomic data. Earlier topologies in Wang et al. (2015) separated *Sampaiozyma* and *Leucosporidium* from the Microbotryales, while *Heterogastridium* and *Reniforma* were placed as sister taxa with low branch support. Li et al. (2020) and Jiang et al. (2024) inferred a topology similar to the present grouping, albeit with low support, and still including *Reniforma* as sister to *Yuzhangia*. The Sporidiobolales (*Sporobolomyces, Rhodotorula, and Rhodosporidiobolus*) is placed in our phylogeny with *Curvibasidium*, *Pseudoleucosporium*, *Nakaseozyma*, *Baiomyces*, and *Fengyania* with a high degree of support, which may also suggest either re-circumscription of Sporidiobolales or the need for additional family- and/or order-level taxonomic description for these genera.

The improved support for the Microbotryales clade and the nearby placement of *Leucosporidium* and *Sampaiozyma* may partly reflect the exclusion of *Reniforma strues* from the present analysis. It is possible that the long branches on which *Yuzhangia* is placed attracted the highly diverged *R. strues* to this clade in earlier analyses (Wang et al. 2015; Li et al. 2020; Jiang et al. 2024). Through sequencing the genome of *R. strues*, we confirmed that despite the loci in public databases being correct, the various protein-coding loci show best BLASTn matches to taxa in a variety of fungal phyla. This suggests that the phylum-level placement of *R. strues* is uncertain, and will likely require additional genome-scale phylogenetic inference to resolve its position. For this reason, *R. strues* was excluded from our analysis.

Well-supported nodes suggest the description of *Aimea* as a new genus including three species (*A. erigeronia*, *A. cardamina*, and *A. sorghi*), which forms a clade with the Microbotryales and *Sampaiozyma*. Because other genera in the class have branch lengths which are either longer or shorter than the branch in common for *Aimea*, this clade is reasonable to describe as a genus. The Microbotryomycetes are well supported as a class, but the nodes along the backbone show short branch lengths and low support, so that many *incertae sedis* classifications remain for family- and order-level clades.

Phylogenetic inference using 7 loci support the distinction of *Aimea* as a novel and distinct genus. The relatively low ANI between isolates (∼70%), and distinct metabolic and slight morphological traits, suggest that these isolates represent unique species under the phylogenetic and morphological species concepts. Attempts to induce mating among the isolates were unsuccessful so far, but fungal sexual cycles can be difficult to recapitulate *in vitro*, even among isolates of the same species (Fraser et al. 2003; O’Gorman et al. 2009).

Given our analysis of the mating loci, these isolates appear to be heterothallic, with *a1* and *a2* alleles at the *P/R* locus and a comparatively conserved *HD* locus. The greater structural variation observed in the *P/R* region, including differences in pheromone gene copy number, is consistent with the broader tendency of the *P/R* locus to be more evolutionarily labile than the *HD* locus in basidiomycetes. In this context, it is notable that recent work in *Cryptococcus* linked pheromone gene duplication to structural instability and expansion of the *P/R* region, likely through the generation of inversion-prone architectures (Coelho et al. 2025). In *Aimea*, variation in pheromone gene number and organization may likewise reflect a dynamic history of local expansion and contraction within the *P/R* locus. This possibility may also be relevant to other Microbotryomycetes, such as *Leucosporidium scottii* (Maia et al. 2015) and Sporidiobolales (Sen et al. 2019; Liu et al. 2025), which exhibit a broadly similar locus organization.

The comparative genomics approach to examine genome annotations appears to reflect an increased abundance of retrotransposons in *Aimea*, given the enrichment of integrase, reverse transcriptase, ribonuclease H, chromo domain, and polyprotein elements (Muszewska et al. 2011) in this genus. However, an important caveat is that the detection of certain functional annotations may be affected by differences in genome sequencing strategy, assembly quality, and gene model prediction across the panel of compared genomes and their respective proteomes. This may be especially relevant for repetitive elements, such as retrotransposon-derived sequences, which are more challenging to detect in fragmented short-read assemblies (Zhou et al. 2020)

Accordingly we cannot exclude the possibility that the lower fragmentation of the *Aimea* genomes relative to other available assemblies contributed to the more frequent annotation of such elements. However, many of the other significantly enriched or depleted annotations are not similarly repetitive and thus are less likely to be biased by the quality of the predicted proteome.

Beyond these retrotransposon-associated features, the proteomes of *Aimea* spp. are depleted of ribonuclease inhibitors (RIs or RNIs) and RNI-like proteins. While these are not known in the literature to be associated with retrotransposon defense, their anti-ribonuclease activities (Kobe and Deisenhofer 1996) could suggest an interaction with retroviral ribonucleases. Proline-rich extensins, also depleted in *Aimea*, are proteins typically known from plant cell walls (Marcus et al. 1991; Saha et al. 2013). However, this same protein family (PR01217) is present in fungal proteins such as the RNA polymerase II RPO21 in *Saccharomyces cerevisiae* (Blum et al. 2025). Due to the presence of this annotated family in proteins of other known functions, the importance of its differential abundance is uncertain. This similarly applies to proteins containing F-box-like and MYND finger domains. ATP-citrate synthase/lyase was notably the only energy metabolism pathway annotation enriched in one of the *Aimea* species. This enzyme is involved in lipid production (Chypre et al. 2012), and in the oleaginous yeast *Yarrowia lipolytica* its heterologous expression results in enhanced lipid accumulation (Zhang et al. 2014). While we have not examined the macromolecular composition of these yeast isolates, the relative enrichment of this annotation suggests that *A. cardamina* may favor lipid accumulation compared to the other species.

Since Ruinen (1961) described the phyllosphere as “an ecologically neglected milieu” sixty-five years ago, the surfaces of plants, especially leaves, have been intensely studied with tens of thousands of metagenomic samples collected and deposited into GenBank (Sohrabi et al. 2023). Despite this breadth of sampling, *Aimea* spp. are not represented in sequence databases including GenBank and GlobalFungi (Větrovský et al. 2020) outside of the isolates presented here. However this appears not to be unusual for the Microbotryomycetes, for which many species and genera are represented by a single isolate (Li et al. 2020; Jiang et al. 2024).

The demonstrated transformability of this genus is relatively unusual in the literature. Of all basidiomycete yeasts, a handful of taxa have published protocols for transformation including the human pathogens in *Cryptococcus* s.s. (Toffaletti et al. 1993) and the closely related yeast-like *Tremella* (Guo et al. 2008), the red yeasts in Sporidiobolales (Takahashi et al. 2014; Pi et al. 2018), and the plant pathogens in Ustilaginaceae including *Ustilago* (Wang et al. 1988) and *Pseudozyma* (Marchand et al. 2007). Given the potential of many basidiomycete yeasts in industrial (Pi et al. 2018) and agricultural (Filonow 1998) uses, as well as their importance as human pathogens (Kim et al. 2013; Liu et al. 2019), additional genetic tools for understanding their biology are extremely important.

Agricultural productivity poses among the greatest challenges to satisfying the world’s demands for food while preserving a livable planet (Baldos and Hertel 2014). Phyllosphere and rhizosphere microbiota are increasingly studied as an important contributor to plant’s defenses against biotic and abiotic stresses (Bulgarelli et al. 2013; Sohrabi et al. 2023). Our description of these novel yeasts associated with plant surfaces contributes to the conversion of some of the unknown “microbial dark matter” to described organisms with transcriptome-informed annotated genomes, metabolic and physiologic observations, and genetic tools for future experimental study.

## Data availability

The authors affirm that all data necessary for confirming the conclusions of the article are present within the article, figures, and tables. Sequencing and assembly data are accessioned under BioProjects PRJNA892096 and PRJNA1443367. Accessions of loci used for phylogenetic analysis are list in Table S2. Scripts, phylogenetic trees, and other data are deposited at https://github.com/liberjul/Yeast_genomes.

## Acknowledgements

The authors thank Ian Medeiros and Keaton Tremble for assistance with genome annotations and phylogenetic inference. We also thank Gregory Bonito for sharing the isolate NB124-2. We thank the curators at the USDA and CBS culture collections for the deposition of strains. We appreciate the feedback on the manuscript provided by Joseph Heitman, Rytas Vilgalys, and Teun Boekhout. J.A.L. received funding support from Duke University, the Duke Microbiome Center, and the Triangle Center for Evolutionary Medicine. S.Y.H. is an Investigator at Howard Hughes Medical Institute. This study was also supported by the National Institute of Allergy and Infectious Diseases of the National Institutes of Health under award R01 AI050113-20 (M.A.C.).

## Figure and Table Legends

**Figure S1.**
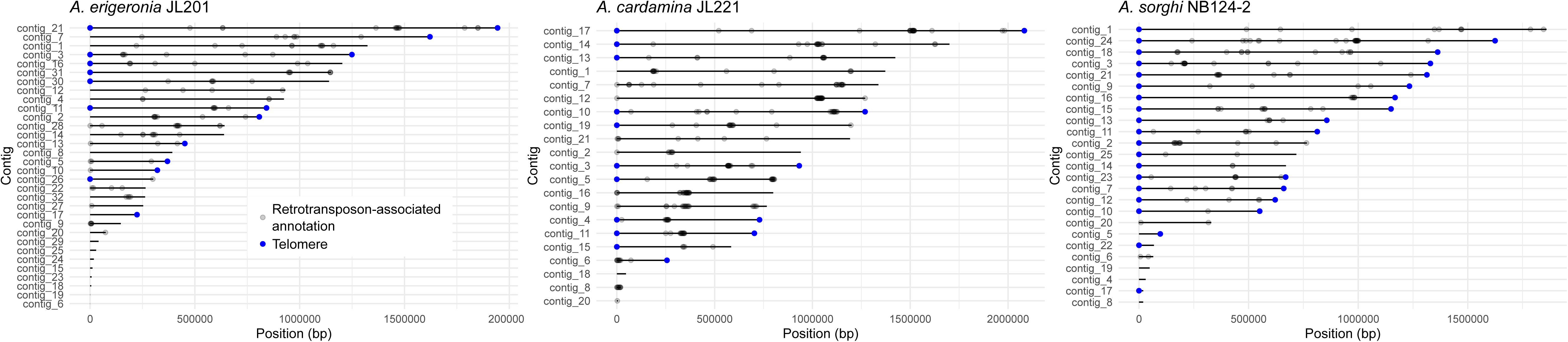
Telomere and putative retrotransposon element positions. Detected telomeric repeats are annotated in blue, and genes with retrotransposon-associated annotations found to be enriched in *Aimea* are annotated with translucent black points.

**Table S1.**
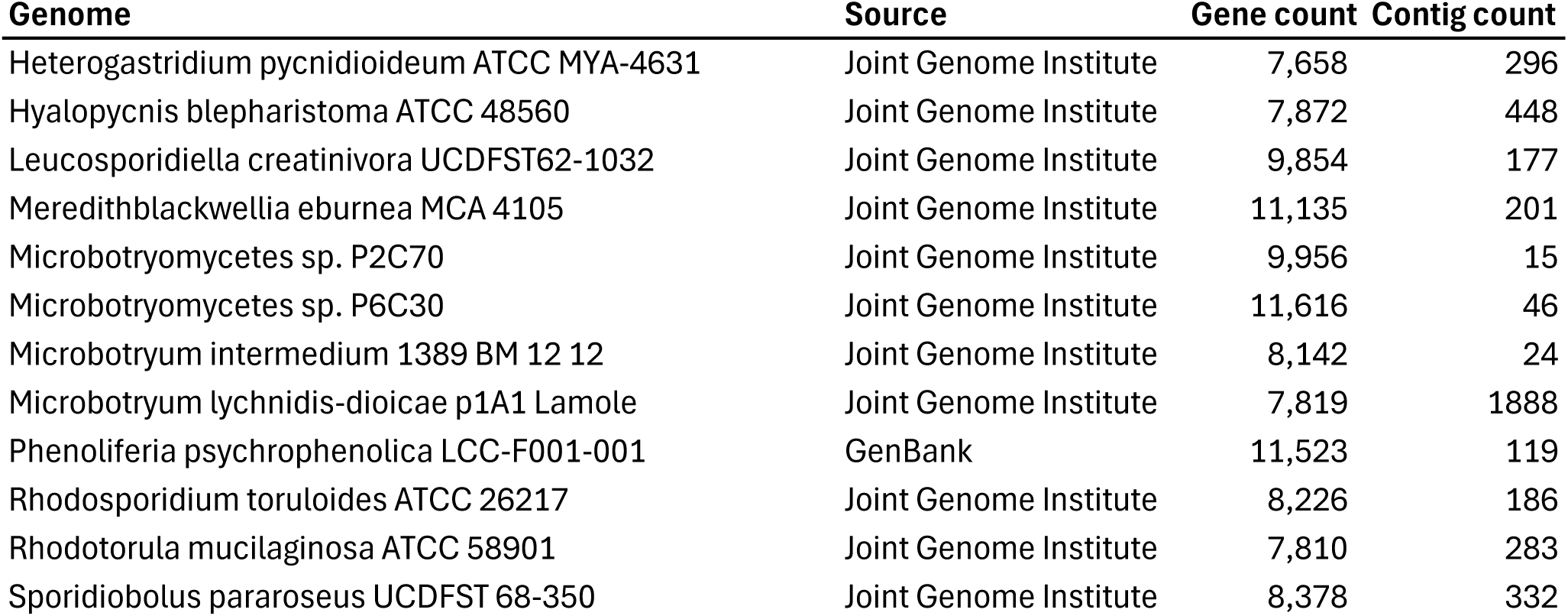
Genomes used for comparative genomics analysis.

**Table S2.**
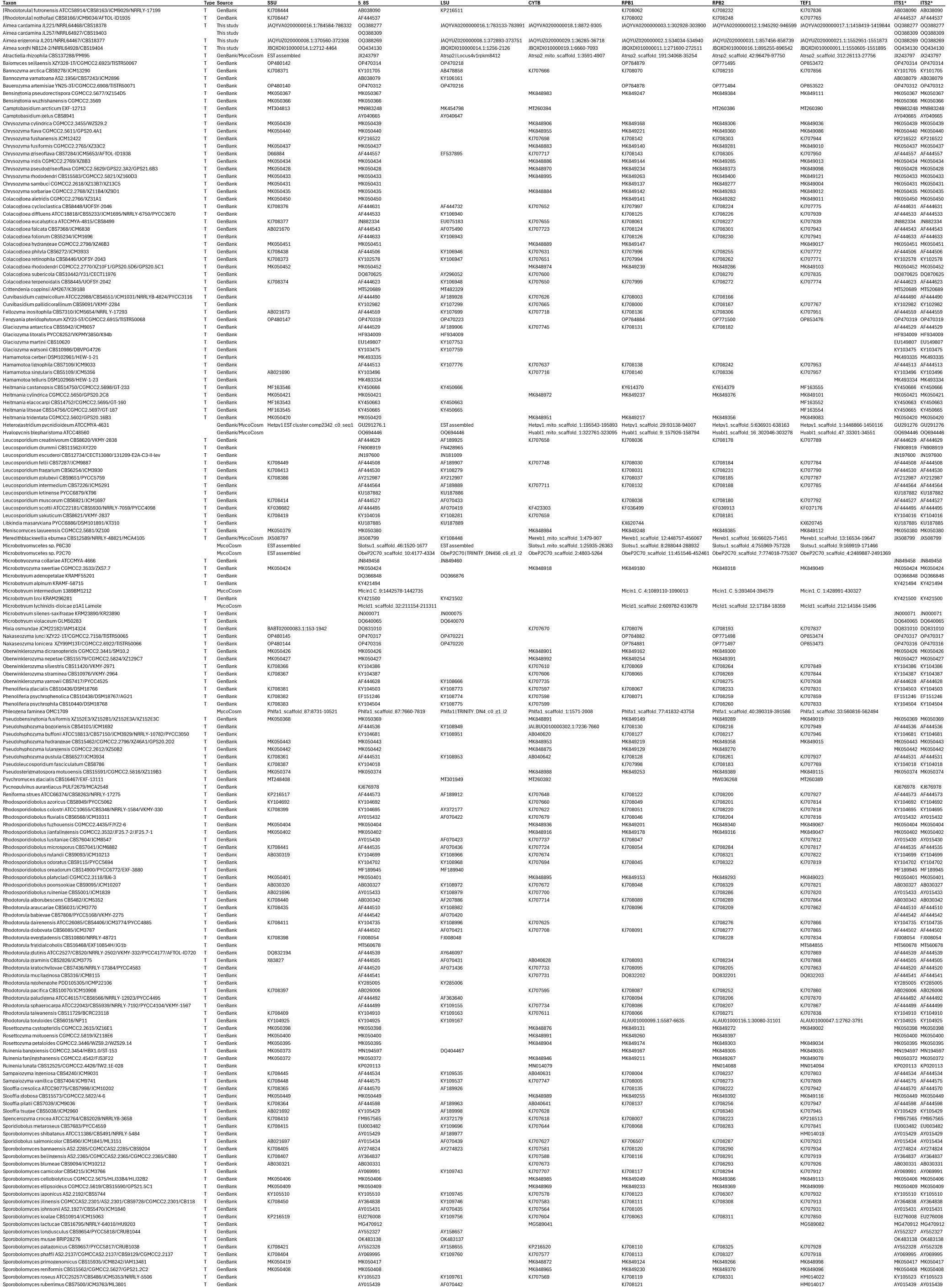
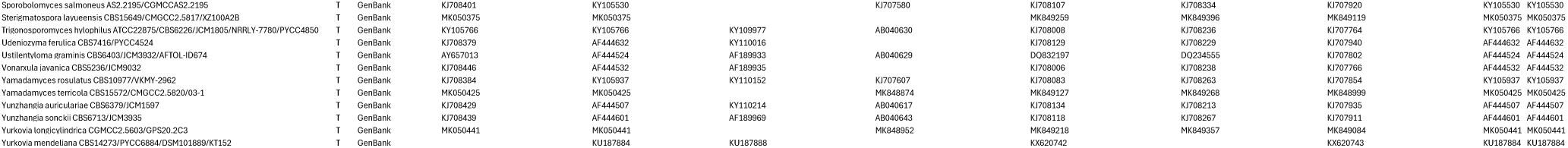
Accessions of data used in phylogenetic inference.

**Table S3.**
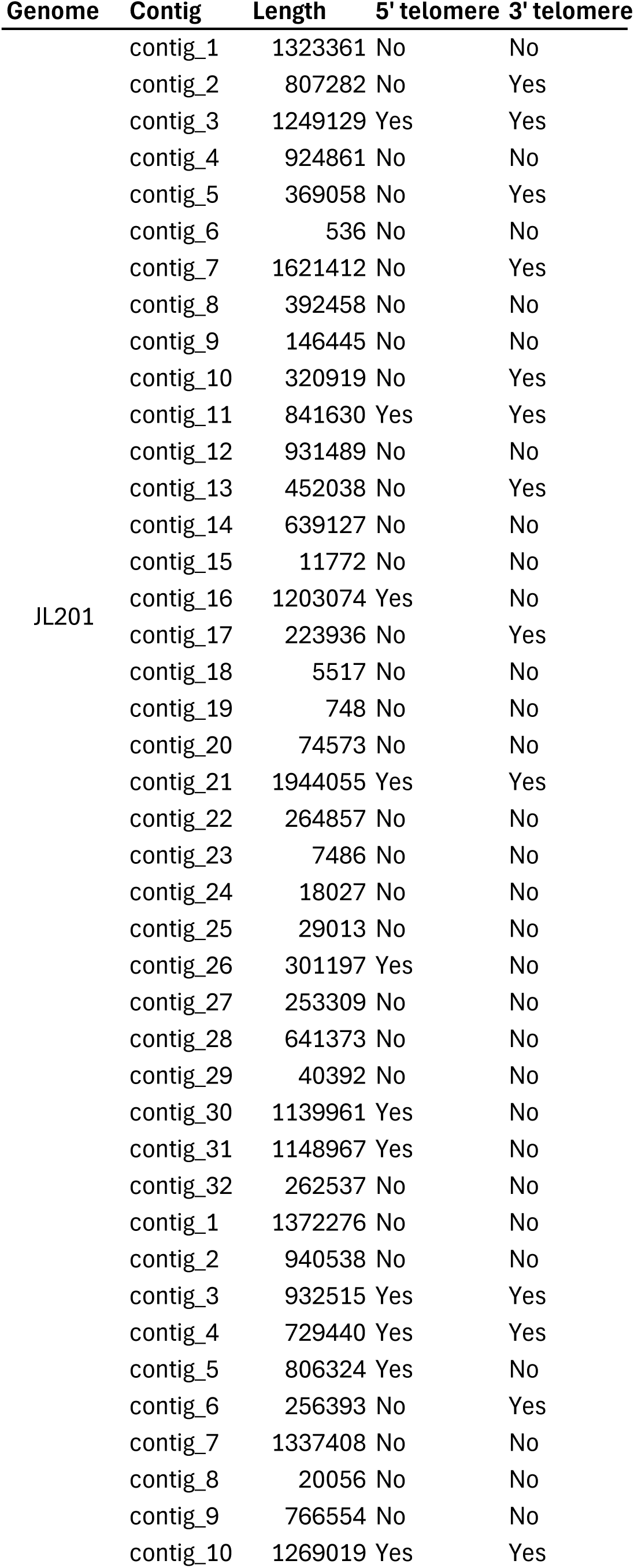
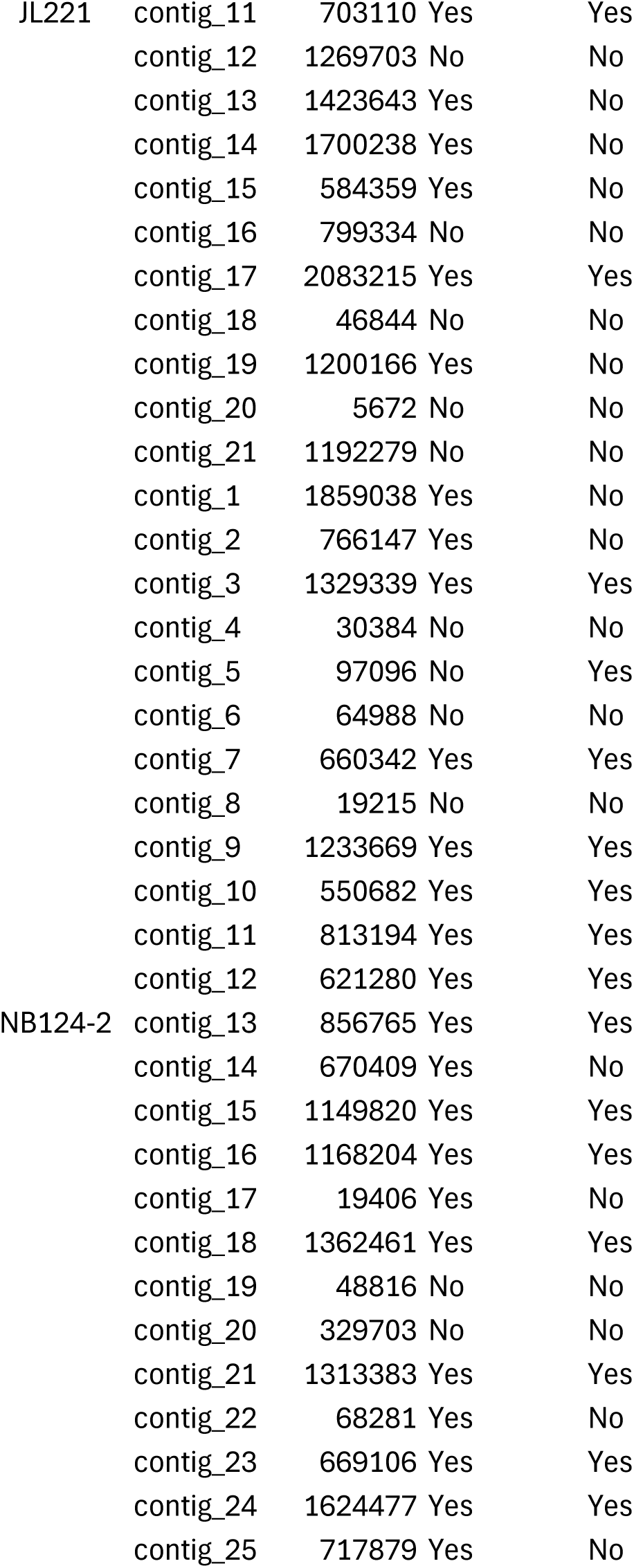
Contig sizes and telomeric repeats.

**Table S4.** Average nucleotide identity between *Aimea* spp. genomes. ANI values are calculated with OrthoANI between the query genome (row) and the subject (column).

|  | **JL201** | **JL221** | **NB124-2** |
| --- | --- | --- | --- |
| JL201 | 100 | 69.1343 | 72.4358 |
| JL221 | 69.1344 | 100 | 69.463 |
| NB124-2 | 72.4368 | 69.4658 | 100 |

**Table S5.** Metabolic and stress tolerance traits. Test numbers come from the deposit form for the Westerdijk Institute’s CBS collection. For assimilation tests, “+” is defined as >50% the optical density of the D-glucose positive control, while “w” is defined as greater than 20% and less than or equal to 50% of the positive control. Other test criteria are defined according to Suh et al. (2008 Nov)

| Test | Substrate | Test Type | JL201 | JL221 | NB124-2 | JL257 |
| --- | --- | --- | --- | --- | --- | --- |
| F1 | D-Glucose | Fermentation | - | - | - | - |
| F2 | D-Galactose | Fermentation | - | - | ? | ? |
| F4 | Sucrose | Fermentation | - | - | ? | ? |
| F5 | Lactose | Fermentation | - | - | ? | ? |
| F11 | Raffinose | Fermentation | - | - | ? | ? |
| F14 | D-Xylose | Fermentation | - | - | ? | ? |
| C1 | D-Glucose | Assimilation | + | + | + | + |
| C2 | D-Galactose | Assimilation | + | w | w | + |
| C3 | L-Sorbose | Assimilation | + | w | w | + |
| C4 | D-Glucosamine | Assimilation | + | w | - | + |
| C5 | D-Ribose | Assimilation | - | - | - | - |
| C6 | D-Xylose | Assimilation | w | - | - | - |
| C7 | L-Arabinose | Assimilation | w | - | - | w |
| C8 | D-Arabinose | Assimilation | w | w | - | w |
| C9 | L-Rhamnose | Assimilation | - | w | - | + |
| C10 | Sucrose | Assimilation | + | + | + | + |
| C11 | Maltose | Assimilation | + | + | + | + |
| C12 | $\alpha,\alpha$-Trehalose | Assimilation | + | + | + | + |
| C13 | Methyl-α-D-Glucoside | Assimilation | + | + | w | + |
| C14 | Cellobiose | Assimilation | - | - | - | - |
| C15 | Salicin | Assimilation | w | + | + | w |
| C16 | Arbutin | Assimilation | w | - | - | + |
| C17 | Melibiose | Assimilation | - | - | - | - |
| C18 | Lactose | Assimilation | - | - | ? | ? |
| C19 | Raffinose | Assimilation | + | + | w | + |
| C20 | Melezitose | Assimilation | + | + | w | + |
| C21 | Inulin | Assimilation | - | - | - | - |
| C22 | Starch | Assimilation | + | + | + | + |
| C23 | Glycerol | Assimilation | + | w | w | w |
| C24 | Erythritol | Assimilation | - | - | - | - |
| C26 | Xylitol | Assimilation | - | - | - | - |
| C29 | D-Mannitol | Assimilation | + | - | w | w |
| C31 | myo-Inositol | Assimilation | - | - | ? | ? |
| C33 | 2-Keto-D-Gluconate | Assimilation | + | w | w | + |
| C34 | 5-Keto-D-Gluconate | Assimilation | ? | ? | ? | ? |
| C35 | D-Gluconate | Assimilation | w | w | - | + |
| C38 | DL-Lactate | Assimilation | w | w | w | w |
| C40 | Citrate | Assimilation | + | + | + | + |
| C41 | Methanol | Assimilation | - | - | - | - |
| C42 | Ethanol | Assimilation | - | - | - | - |
| C48 | Palatinose | Assimilation | + | w | w | + |
| C56 | Gentiobiose | Assimilation | w | - | - | w |
| N4 | L-Lysine HCl | Assimilation | ++ | ++ | + | ++ |
| N5 | Cadaverine HCl | Assimilation | ++ | + | ++ | ++ |
| N6 | Creatine | Assimilation | - | - | - | - |
| N7 | Creatinine | Assimilation | - | - | - | - |
| N8 | Glucosamine | Assimilation | ++ | +++ | - | - |
| N9 | Imidazole | Assimilation | - | - | - | - |
| N10 | D-Tryptophan | Assimilation | + | + | - | - |
| V1 | w/o vitamins | Vitamin | + | - | - | + |
| V2 | w/o myo-Inositol | Vitamin | + | + | + | + |
| V3 | w/o Pantothenate | Vitamin | + | + | + | + |
| V4 | w/o Biotin | Vitamin | + | + | + | + |
| V5 | w/o Thiamin | Vitamin | + | - | - | + |
| V6 | w/o Biotin & Thiamin | Vitamin | + | - | - | + |
| V7 | w/o Pyridoxine | Vitamin | + | + | + | + |
| V8 | w/o Pyridoxine & Thiamin | Vitamin | + | - | W | + |
| V9 | w/o Niacin | Vitamin | + | + | + | + |
| V10 | w/o PABA | Vitamin | + | + | + | + |
| O1 | Cycloheximide 0.0001 | Resistance | + | W | + | + |
| O2 | Cycloheximide 0.001 | Resistance | - | - | - | - |
| O3 | Acetic acid 0.01 | Resistance | - | - | - | - |
| O4 | 50% w/v D-Glucose | Resistance | W | + | W | W |
| O5 | 60% w/v D-Glucose | Resistance | - | - | - | - |
| O6 | 10% w/v NaCl | Resistance | + | + | + | + |
| O7 | 16% w/v NaCl | Resistance | - | - | - | - |
| O8 | Growth at pH=3 | Resistance | W | W | ? | ? |
| O9 | Growth at pH=9 | Resistance | - | - | ? | ? |
| O10 | Fluconazole | Resistance | ? | ? | ? | ? |
| T1 | Growth at 25C | Temperature | + | + | + | + |
| T2 | Growth at 30C | Temperature | + | + | + | + |
| T3 | Growth at 35C | Temperature | - | - | - | - |
|  | Urease activity | Enzyme | + | + | + | + |
|  | Starch formation | Product | - | - | - | - |
|  | Acetic acid formation | Product | - | - | - | - |
|  | Gelatin liquefaction | Enzyme | W | + | + | + |

